# Video-recorded measures of blinking differ with group size in red deer *Cervus elaphus* and fallow deer *Dama dama*

**DOI:** 10.1101/2024.01.02.573958

**Authors:** Thomas R. Howard, Sean A. Rands

## Abstract

The group vigilance hypothesis suggests animals within a group have a lower overall vigilance level than solitary individuals due to the collective responsibility of predator detection. Previous methods of studying vigilance have used head position as a proxy to indicate a state of vigilance, but it has been suggested that this method is too ambiguous, and the observer cannot accurately ascertain the exact behavioural state of the focal animal. Here, we consider blinking as a proxy for vigilance, as the eye-closure during a blink momentarily reduces the animal’s information about the environment, increasing the risk of unanticipated predator attack. We videoed semi-captive herds of red deer *Cervus elaphus* and fallow deer *Dama dama*, and quantified their blinking behaviour using behavioural logging software. We show that blink rate decreased with increasing group size in both species, suggesting that blinking may be tied with social vigilance rather than anti-predator vigilance. Similarly, the length of the inter-blink interval increased with increasing group size in red deer. We discuss the suitability of blinking measures for assessing vigilance behaviour.

## INTRODUCTION

Forming groups with other individuals allows members of many prey species to reduce their predation risk. By grouping together, a larger group has a higher chance of detecting an attack before it occurs, and if an attack does occur, an individual’s risk is diluted by being in a group compared to if the individual was solitary (Allan and Hill, 2018; Bednekoff and Lima, 1998; Lehtonen and Jaatinen, 2016). Advantages from group-living also come from combined anti-predator behaviours of all the group’s members, such as scanning behaviour, which reduces predation risk as it increases the likelihood of predator detection (Beauchamp, 2019; Childress and Lung, 2003). Within the group, there are more individuals scanning the surrounding environment, so when coupled with diluting individual risk of being captured, group members can reduce overall vigilance with little cost (Sansom et al., 2008). Lone individuals are responsible for all of their vigilance duties, and therefore may have to forego other activities such as foraging or mating to reduce its risk of being attacked, for instance, bison *Bison bison* and wapiti *Cervus canadensis* show a reduction in food intake when vigilance increased (Fortin et al., 2004). In larger groups, vigilance responsibilities may be split between the members of the group, and therefore, each individual is required to dedicate significantly less time to vigilance (Beauchamp, 2019). For example, there is an observed negative relationship between group size and individual vigilance and a positive relationship between group size and overall herd vigilance in Tibetan gazelle *Procapra picticaudata* (Li and Jiang, 2008). This pattern is described by the ‘Group Vigilance Hypothesis’ or ‘the many eyes effect’ (Allan and Hill, 2018; Beauchamp, 2019; Sansom et al., 2008).

By having to reallocate and redistribute time from foraging to being vigilant, vigilance is therefore costly (Blanchard et al., 2017). Due to this, when perceived risk of predation is proportionally higher, the allocation of time to vigilance will increase (Eccard et al., 2017; Yorzinski et al., 2021). Referred to as the ‘Risk Allocation Hypothesis’, this pattern has been observed both spatially, where animals may avoid a specific high risk area, and temporally, where they only forage during certain lower-risk time periods throughout the day (Creel et al., 2008). An individual in a smaller group is more at risk than an individual in a larger group, due to the higher chance of being predated, and therefore individuals in smaller groups are expected to be more vigilant (Lehtonen and Jaatinen, 2016; Rowe et al., 2023; Sansom et al., 2008). The costs and risks associated with vigilance mean that it is a potentially useful measure of welfare (Cherry et al., 2020; Flamand et al., 2025; Lelláková et al., 2023; Tomberg et al., 2024) and affective state (Guy Beauchamp, 2017; Crump et al., 2018; Welp et al., 2004), particularly because it is a non-invasive measure that can be observed without manipulation.

However, vigilance is tricky to measure, as it can occur when the animal is conducting a lot of different behaviours, and the effort and quality of the vigilance conducted may depend upon the current behaviour, the internal state of the animal, and the social and physical environment in which it is found. Commonly, vigilance is measured using proxy behaviours, most commonly using the position of the animal’s head as a cue for vigilance. However, there is a lot of variation within the literature, with different authors defining vigilance with different, and often hard to quantify, behaviours leading to varied results (Allan and Hill, 2018; Fernández-Juricic and Beauchamp, 2008; Pecorella et al., 2019; Yorzinski et al., 2021). Recently, blinking has been explored as an alternative proxy of vigilance effort. The term ‘blinking’ refers to eye-blinking, the period of time in which visual input is periodically, and temporarily, obstructed due to eye lid closure – during this time, visual vigilance cannot occur (Rowe et al., 2023). Blinking is necessary in order to maintain ocular health and therefore benefits the individual, but it is also costly due to the inhibition of vigilance behaviours. Therefore, it is expected that blinking should be strategically adjusted so that in high-risk environments, or where predation risk is high, blinking should be repressed to reduce the loss of visual input (Cross et al., 2013; Merkies et al., 2019; Ranti et al., 2020; Yorzinski et al., 2021). While seemingly only a small loss of time, the additive effect of blinking can lead to a large proportion of an individual’s visual input being blocked. Blinking has been studied in humans, showing that individuals will blink less often when faced with a cognitively demanding task (Hoppe et al., 2018) or when watching content that is perceived to be important (Ranti et al., 2020). These same principles can be applied to vigilance in other species, whereby an individual may supress blinking behaviour when group size is smaller due to the increase in vigilance demand (Rands, 2021). This suppression may occur as a reduction in blink duration, or an increase in the interval between blinks so that the blink rate is reduced (Matsumoto-Oda et al., 2018; Yorzinski, 2016).

This study tests how group size affects key components of blinking behaviour (blink rate, blink interval and blink duration) between fallow *Dama dama* and red deer *Cervus elaphus*. An earlier study on red deer (Rowe et al., 2023) concluded that blinking was positively correlated with group size, but the study was conducted by visual observation of blinks in the field. Because blinks are fast behaviours that take a fraction of a second, it is feasible that the data recorded were subject to unintentional bias in identifying these fast behaviours. The current study therefore uses video footage of individual behaviour, which should provide accurate, unbiased data. Previous studies in other species (G. Beauchamp, 2017; Matsumoto-Oda et al., 2018; Rands, 2021; Tada et al., 2013; Yorzinski, 2020; Yorzinski and Argubright, 2019) have shown that video analysis is an accurate means of collecting this information. We would expect that as group size varies, both deer species should modify their blink rate, duration, and length of time between blinks – individuals in a larger group would be expected to be less vigilant, therefore blinking more frequently and for longer durations with shorter inter-blink intervals, as predicted by the group vigilance and risk allocation hypotheses (Beauchamp, 2019; Eccard et al., 2017; Li et al., 2009).

## METHODS

### Study site and subjects

The study was carried out between two sites in Ashton Court Estate, Bristol, England (51.4440°N, 2.6378°W). Ashton Court Estate is a publicly-accessible country estate that covers 850 hectares of grassland and woodland, within which there are two enclosures for red and fallow deer, both of which are approximately 40 hectares in area (enclosures are separated to prevent mixing), and which have previously been described in (Hoyle et al., 2021; Rands et al., 2014; Rowe et al., 2023). The red deer park at Ashton Court Estate comprises of largely open grasslands, with various wooded areas scattered around their habitat. The herd consists of *c*.130 individuals, which is made up of adult and juvenile, male and females, all of which can freely mix with one another. The fallow deer herd consists of *c*.65 individuals, which reside in a more covered enclosure, with approximately half of the enclosure covered by woodlands. The fallow deer herd also consists of adult and juvenile male and females which could also mix freely. Both herds are managed by Bristol City Council. Although the deer are kept in fenced enclosures, they have constant contact with humans and dogs, who can walk around and through their space. Similarly, motor vehicles, bicycles and frequent hot air balloons pass by or over their space.

Ethical approval to conduct the observations conducted in this study was given by the University of Bristol Animal Welfare Ethical Review Board (University Investigation Number UB/21/004).

### Focal observations and data collection

Behavioural observations were carried out between February and May 2022, between 10:00 and 14:00. Individuals were filmed and recorded using a Nikon D3300 camera mounted to an Opticron ES 80 GA ED scope *via* a T2-mount and Opticron Photoadapter; the position of the observer relative to the herd varied according to herd position, but was approximately 100 m. Videos were recorded in mp4 format at 1080p resolution and at 50 frames per second. Individuals were recorded for five-minute periods, which were subsequently broken down into one-minute intervals. At the beginning of a study day, no observations were made within ten minutes of observer arrival to allow for habituation. If any major disturbance occurred, then an observation was halted and restarted following a 10-minute break once the disturbance had finished. This halting of observation only occurred once during the study period.

Individuals were randomly selected by using total herd size, inputting this as an upper bound into a random number generator and then using the randomly generator number to count in from either left-most individual or right-most, depending on a coin flip. Upon observation, individuals were sexed using visually indefinable features of morphology (such as body size and presence of antlers). If an individual was observed to be suckling, the individual was assumed to be a juvenile and discarded, due to potentially abnormal behaviour that might be observed in this age class, whereby it is likely calves are not independently acting as individuals, and heavily relying upon cues from mothers (de la Peña et al., 2021; Pecorella et al., 2019; Rowe et al., 2023); if this occurred, the randomisation selection processes was restarted and another individual was chosen. Prior to observations, a total herd size count was conducted, and after an individual had been randomly sampled, both before and after its observational period, the local group size (for that individual) was also counted. Group size was defined using nearest neighbour clusters, and distances of five body lengths between individuals, as discussed by (Rands, 2015) for group associations. It was also ensured that the distance between observer and focal individual was no less than 50m, (measured using Nikon Forestry Pro II Laser Rangefinder).

Our initial dataset contained observations of both male and female individuals. However, because there were relatively few antlered males observed in the herds (7 red deer males and 5 fallow deer males), the random sampling, coupled with visibility and the movement patterns and tendency of the males to aggregate in small single-sex groups, tended to oversample these antlered individuals. Their behaviour was noticeably different to those of the females, and we decided to drop these data from the study. The supplementary dataset (Howard and Rands, 2025) includes these male data, but we do not consider them further here.

### Behavioural and statistical analysis

In order to record state and point behaviours, video files were exported into BORIS, a free behavioural logging software (Friard and Gamba, 2016). Behaviours were defined as ‘blinking’ (rapid and full eyelid closure, as shown in Figure 1) and ‘blink duration’ (duration, of which eyelids are closed). Furthermore, the duration of how long an individual maintained its head position was also recorded; ‘up’ (head perpendicular to the ground, or at an upward angle) or ‘down’ (head angled towards the ground, less than perpendicular to the ground). Video footage was uploaded and played back; once a blink was recorded as starting, its duration was measured by counting frames using the frame-by-frame mode in BORIS.

**Figure 1.**
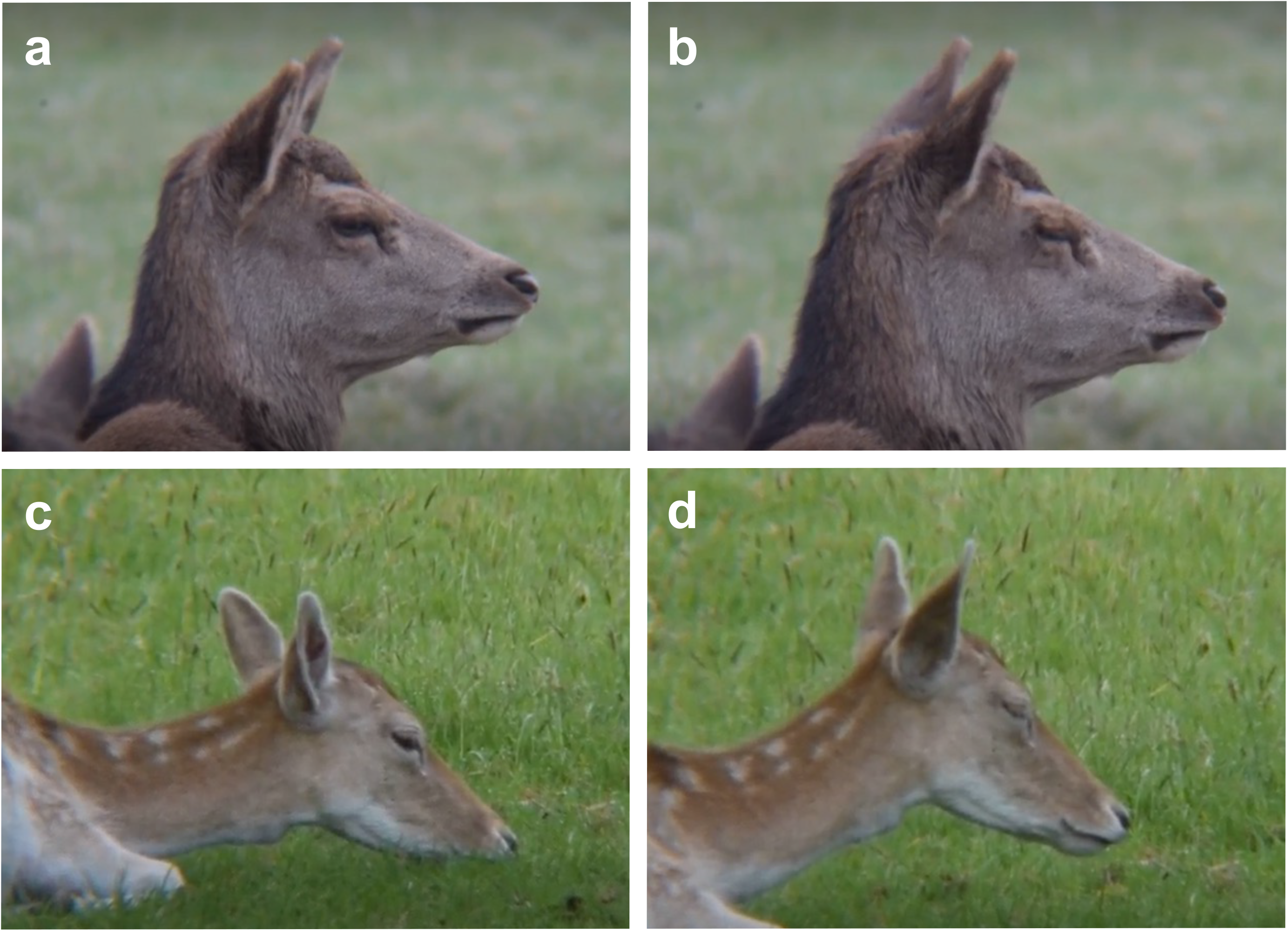
Examples of blinks in red deer (**a**: eyes open immediately before a blink; **b**: eyes closed mid-blink) and fallow deer (**c**: eyes open immediately before a blink; **d**: eyes closed mid-blink).

Each five-minute recording session was broken down into one-minute segments. Upon playback, if the focal individual’s eyes were obscured from view for more than ten seconds, we discarded that one-minute segment. Using the remaining segments, we calculated the blink rate by summing the total number of blinks across the valid recording segments (where a valid recording segment was assumed to occur when a focal individual’s eyes were observed undisturbed) and dividing by the number of valid segments. Blink duration was calculated using the frame-by-frame tool in BORIS. The first frame was counted from the first movement of eyelids, and the final frame was when eyelid movement ceased. Inter-blink interval was calculated as the difference in time between two consecutive blinks. If a segment of recording was discarded due to obstruction of focal individual, we did not consider the length of time between the last blink before and the first blink after the discarded segment, as we did not know whether the individual had blinked during the discarded period.

Statistical analysis was performed using *R* 4.3.0 (R Development Core Team, 2023), using *ggplot2* 3.4.4 (Wickham, 2009) for graphics. All data and code are freely available at (Howard and Rands, 2025). The relationships between group size and the focal individual’s behaviour were considered for mean blinking rate, for mean blinking duration, and for mean interval between blinks for the two species, using a linear model. In order to satisfy model assumptions, we transformed the data for some of the measures, using square roots for the three red deer measures, and the natural logarithm for the fallow deer interval measure (with fallow deer rate and duration remaining untransformed).

The impact of group size was also considered for the proportion of time that the red deer and fallow deer had their heads down. The proportional data had a large number of ‘1’ values, so we used fractional logistic regressions by running a generalised linear model using a quasibinomial distribution with a logit link function.

### Pseudoreplication

Due to the unavoidable chance that individuals within the herd were resampled, there is chance that a degree of pseudoreplication occurred (Hurlbert, 1984). To test whether pseudoreplication could skew the data set, we generated 100,000 datasets for each species and each of the three proxies that we were measuring: blink duration, interval, and blink rate. Each dataset assumed a herd size of 123 females for red deer, and 60 females for fallow deer. Following the procedure described in (Hoyle et al., 2021; Rowe et al., 2023), the observations within each dataset (51 observations for red deer, 36 observations for fallow deer) had a randomised focal identity allocated to them, independent of the identities allocated to the other observations within the dataset. For example, for a red deer observation, that observation could be allocated the identity of any one of the 123 females present; within the same dataset, another female observation would again be allocated with the identity of any one of the 123 females present, and so the same identity could be allocated between observations within the same dataset multiple times by chance alone. Having randomly allocated identities to all of the observations within a simulated dataset, we then filtered these so that if multiple observations were allocated to the same individual, then only one of these observations (randomly chosen from the individual’s set) was used, with the others discarded from the simulated dataset. Having filtered a simulated dataset to remove repeated observations, we then performed linear modelling as described above, and harvested the resulting simulated *p* value. These simulated *p* values were collected for all 100,000 simulations, and the proportion ψ falling below the set significance level (*p* ≤ 0.05) was calculated. If pseudoreplication were not an issue, we would expect to see a higher proportion of simulated significant results for cases where the ‘real’ data were significant, giving a high value of ψ. Alternatively, if simulations demonstrated that statistically significant results were unlikely, we would expect to see a low value of ψ. This means that if we see a significant result with *p* < 0.05 in our analysis of the unmanipulated results, but then find that ψ is low for the corresponding resampled data, then the significant result observed is most likely to be due to pseudoreplication through the repeated sampling of the same individuals who may be showing anomalous behaviour that does not reflect that of other herd members.

## RESULTS

In this study, 110 five-minute observations were made (n = 550 one-minute segments), of which 119 one-minute segments were discarded due to visual obstruction of focal individual. After removing male data (see methods), our usable dataset contained 51 observations of red deer and 36 of fallow deer. Within these, we recorded 1156 individual blinks (red deer *n* = 761, fallow deer *n* = 395). Group size varied between species: red deer showed a mean of 61.08 ± 6.00 (SE) individuals (SD = 42.15), and fallow deer, 21.47 ± 2.88 individuals.

For red deer, blinking rate decreased with group size (*F*_1,49_ = 10.49, *p* = 0.002, Figure 2, *ψ* = 0.983). Blink duration was not related to group size (*F*_1,49_ = 2.50, *p* = 0.121, *ψ* = 0.103). The interval between blinks was related to group size (*F*_1,48_ = 6.99, *p* = 0.011, *ψ* = 0.806, Figure 3). There was no relationship between the proportion of time with the head up and group size of the focal individual (*t*_47_ = 1.086, *p* = 0.283, *ψ* = 0.004). We note that all pseudoreplication test statistics reported in the results align with predicted values, and so any unintended repeated measurements of focal individuals did not unduly influence the statistics reported.

**Figure 2.**
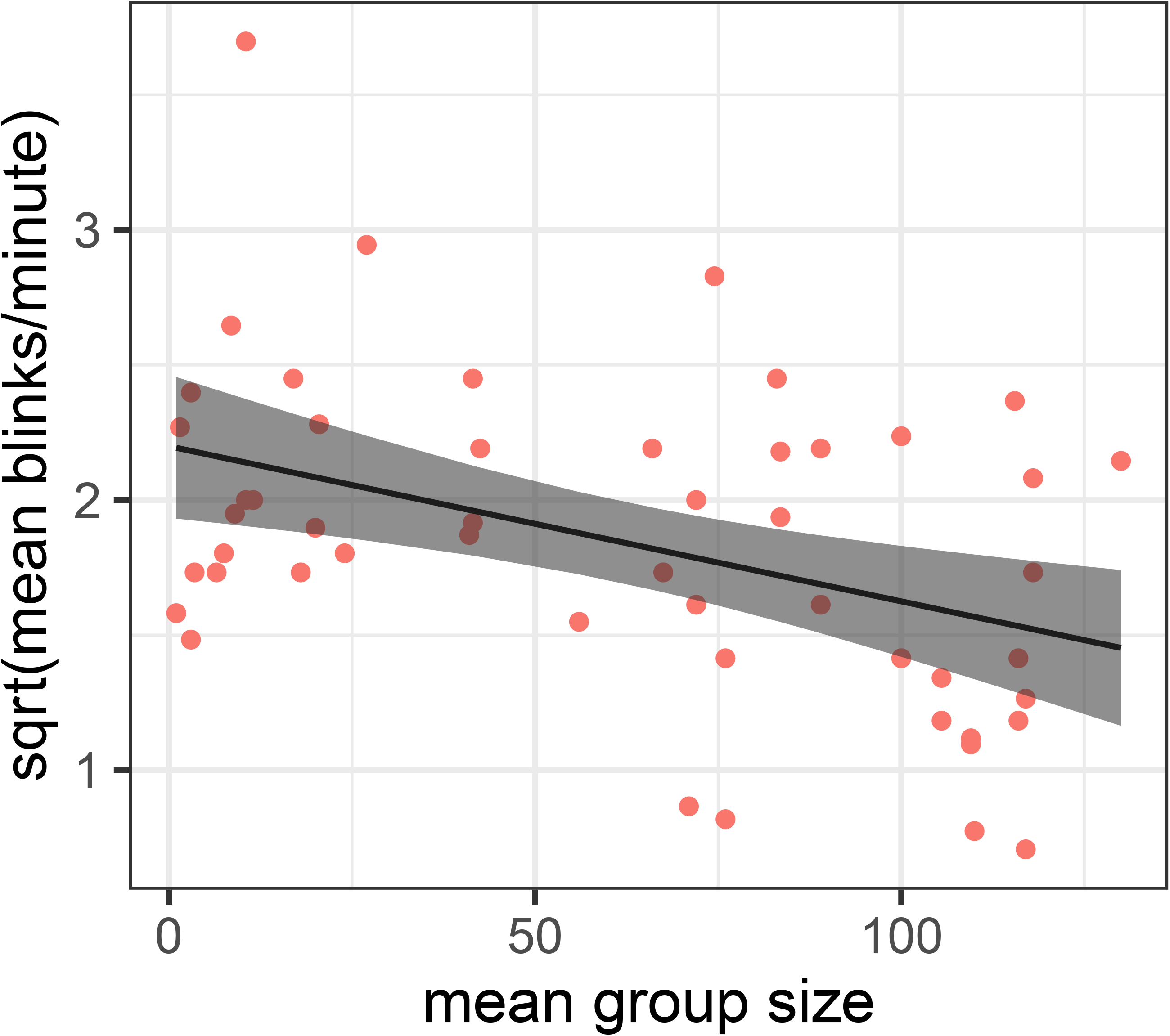
The relationship between mean group size on mean blink rate in female red deer. The figure shows fitted lines (with 95% confidence interval) from the linear model.

**Figure 3.**
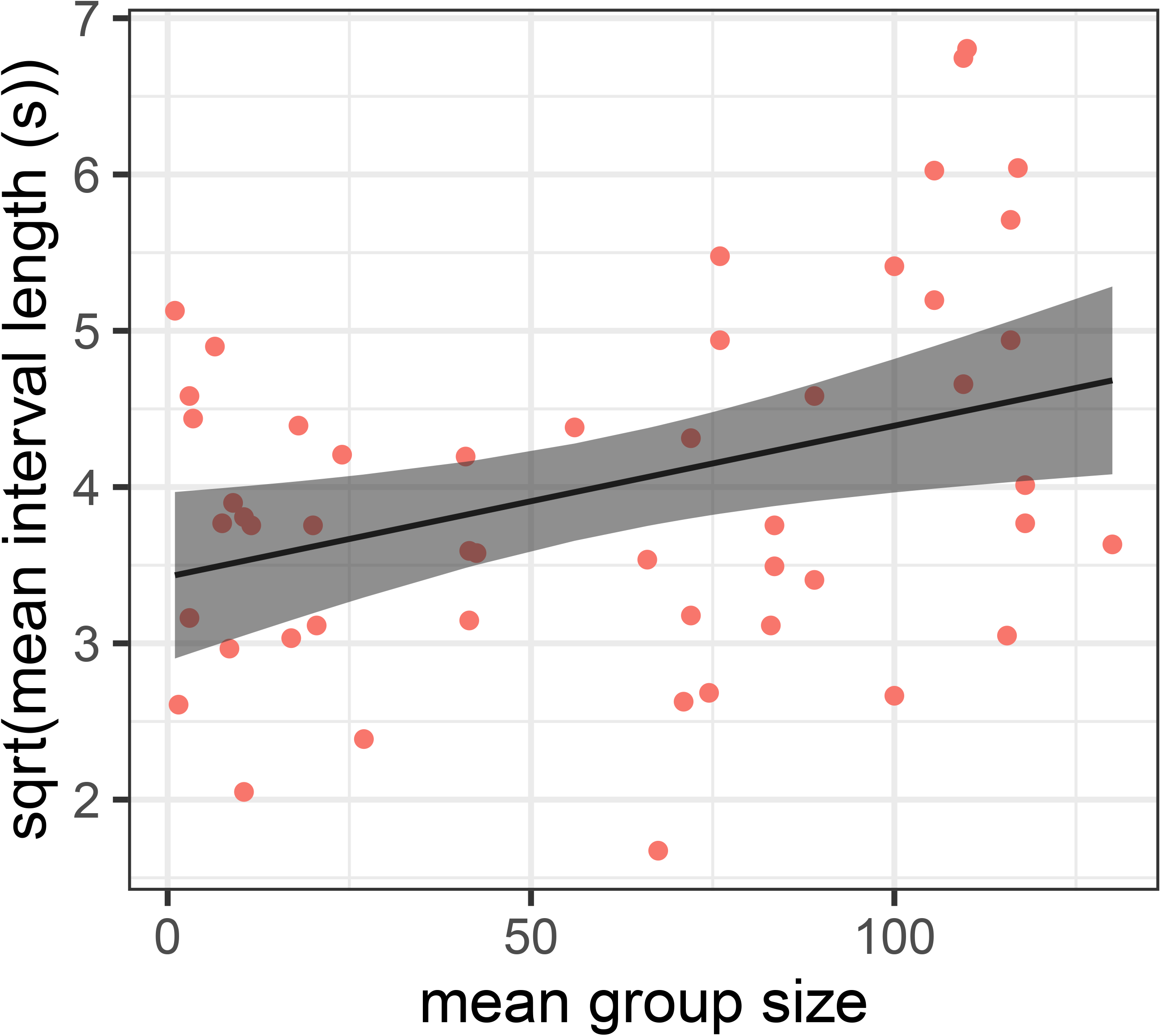
The relationship between mean group size and mean inter-blink interval length in female red deer. The figure shows fitted lines (with 95% confidence interval) from the linear model.

In fallow deer, the blinking rate decreased with group size (*F*_1,34_ = 9.26, *p* = 0.005, *ψ* = 0.878, Figure 4), but group size showed no relationship with either duration (*F*_1,31_ = 0.34, *p* = 0.564, *ψ* < 0.001) or interval (*F*_1,31_ = 2.95, *p* = 0.096, *ψ* = 0.056). Similarly, there was no relationship between the proportion of time with the head up and group size of the focal individual (*t*_34_ = 1.583, *p* = 0.123, *ψ* = 0.085).

**Figure 4.**
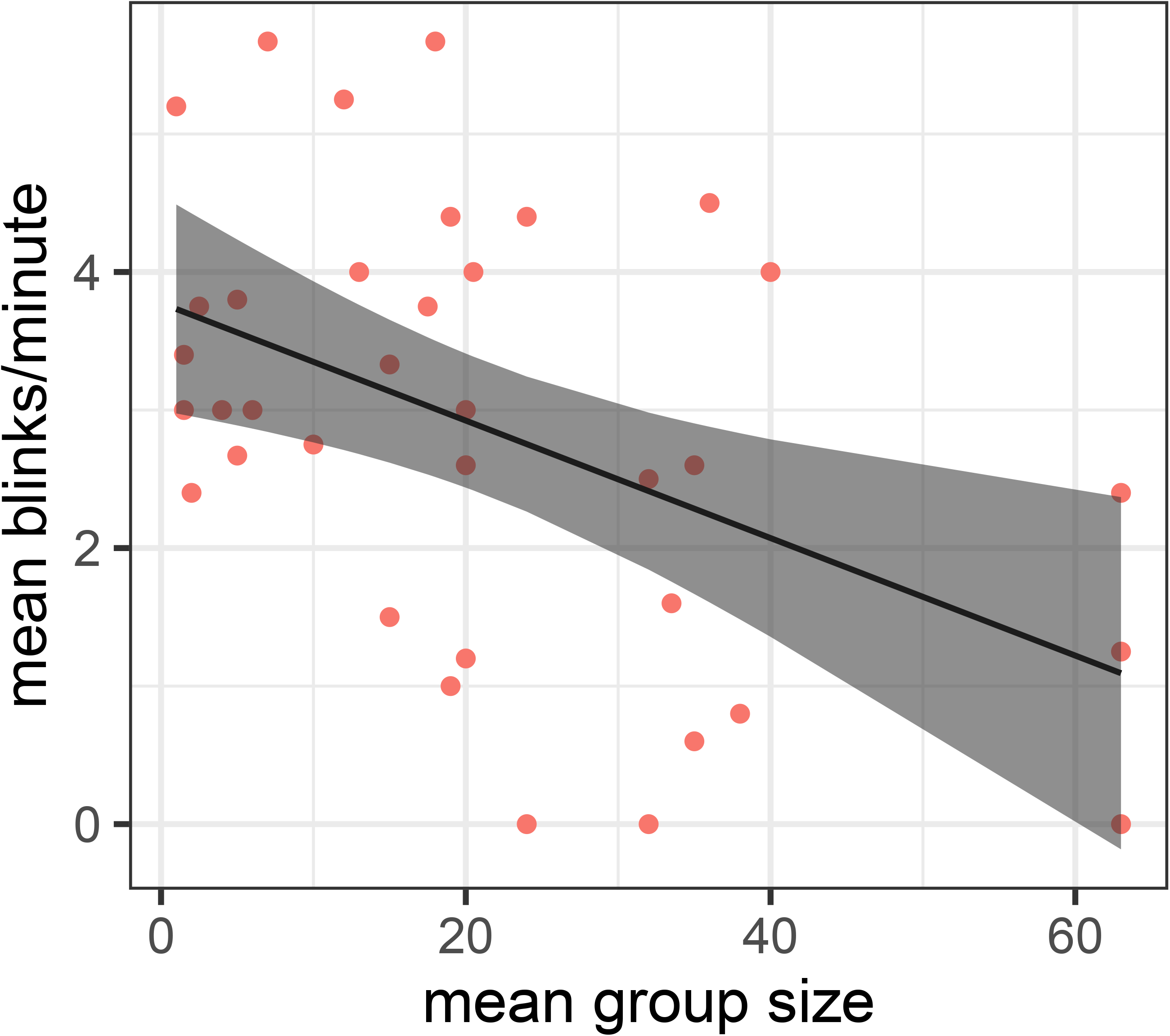
The relationship between mean group size on mean blink rate in female fallow deer. The figure shows fitted lines (with 95% confidence interval) from the linear model.

## DISCUSSION

Following the group vigilance hypothesis, we expected to observe both species of deer showing lower levels of vigilance with increasing group size. Assuming blinking is a good proxy of vigilance, we expected that in larger groups, blink rate and blink duration would be higher, and blink interval would be shorter for the same reasons. However, our results suggest the opposite – blink rate decreased with an increase in group size in both species, although neither of the other measures showed any relationship with group size. The social element suggests that attention by individuals may be being focussed on the social environment rather than predation risk. Social vigilance is likely to occur due to increased competition to food, or during breeding seasons when harassment by males is common (Favreau et al., 2010; Pecorella et al., 2019). Furthermore, individuals may be investing time in scanning the group in order to gain information about both social information and external events (Fernández-Juricic et al., 2005). We observed that blink rate in both species decreased as group size increased. The data for this study was collected between February and May, and as a result, individuals could be displaying residual stress and therefore heightened vigilance as a result of the intense male interactions that occur during the rut (de la Peña et al., 2021; Kjær et al., 2008; Zheng et al., 2013). In mixed-sex groups, similar to the group observed in this study, vigilance was observed to not decrease as group size increased in Pere David’s deer *Elaphurus davidianus* (Zheng et al., 2013), which the authors attributed to the threatening nature of males during the rut. Furthermore, the group of deer studied in (Zheng et al., 2013) were similar to the groups in the current study, in that they resided in a protected reserve and human activity was the biggest threat to them. We could debate whether the deer in the current study are experiencing ‘natural’ predation risk (see the discussion of (Rowe et al., 2023)), but regardless of this, the current study demonstrates that even without the presence of a natural predator, deer are able to adjust vigilance in response to threats that might not necessarily cause loss of life (Pecorella et al., 2016). In another study, (Childress and Lung, 2003) also found group vigilance to increase with herd size. There is a clear inconsistency with group size effect on vigilance within the literature, but this is likely to be due to specific and individual costs and benefits of being vigilant for each herd, which is a possible reason as to why the current study did not find results that matched the expected hypotheses.

As previously mentioned, human disturbance is the leading threat that faces the herds of deer that were used for this study. Red deer have been observed to display the same levels of vigilance in both predator absent and predator present areas (Ginkel et al., 2019). Previous studies have further shown that red deer respond to human disturbance by increasing vigilance (Jayakody et al., 2008). Moreover, the frequent recreational use of deer habitat is shown to elicit behavioural changes by red deer stags (Sibbald et al., 2011). Ashton Court, the study site, is a public area whereby the deer are frequently in contact with both humans and dogs, and so while it could be argued that the deer are habituated to these, repeated exposure to such a stimulus does not necessarily lead to an loss of the reaction (McSweeney and Swindell, 2002). Nevertheless, it would be instructive to explore how the behaviour of these deer compares to those experiencing more apparent predation risk.

The novel part of this study was to develop the quantification of blinking as a proxy measurement of vigilance. While blinking as a proxy has been used in studies to assess vigilance (Matsumoto-Oda et al., 2018; Rowe et al., 2023; Tada et al., 2013), this study was the first to measure all three aspects of blinking (rate, duration, and inter-blink interval) simultaneously in Cervidae, using a video recording device and behaviour logging software in order to maintain a high level of accuracy. Our main finding is that blink rate, while affected by group size, does not increase as expected in red and fallow deer, which contrasts with Rowe et al. (Rowe et al., 2023) who measured blinking in the same herd of red deer. A key difference between the two studies is that Rowe et al. (Rowe et al., 2023) counted blinks in the field, in real time, and therefore human error may have a greater impact on the results, although we note that the studies occurred at different times of year. This is not to say that the current study is without observer bias, but having a video recording of the observations means that the results can be verified by a third party – an improvement that is recommended should this study be repeated. Rowe et al. (Rowe et al., 2023) also only used blink rate as a measurement, whereas the current study also used blink duration and inter-blink interval. Blink duration has not been used to evaluate vigilance in groups of Cervidae before, although we note that Tada et al. (Tada et al., 2013) measured the blink duration across 71 different primates and found, similar to our study, no correlation with group size, which suggests that blink duration might be physiologically constrained. Tada et al. (Tada et al., 2013) further suggest that eye-blinks may be governed not only by external threats but also social factors too, which would explain why blink rate in the current study decreased as group size increased, as individuals were dedicating more visual attention to the activity of their conspecifics.

Evaluating the three potential proxies for vigilance used in this study, blink duration is the measure that provides the least evidence that would allow us to assess vigilance in our study species. This is due to consistent lack of significant results throughout the study and low confidence from pseudoreplication tests. Lack of specialised equipment could be the reason for a lack of identifiable correlation between blink duration and group size. Given that a blink is less than 250ms, using non-specialist videography equipment is unlikely to pick up small changes across long distances. Secondly, blink interval was only found to be significant in red deer, and pseudo-replication tests reinforce the confidence in these results. This needs to picked apart, as the rate of blinking should be inversely related to the length of the intervals between blinks, and it is likely that the mismatch seen is due to the lower numbers of interval measurements taken, which was affected by gaps occurring in the data due to missing one-minute intervals. Finally, blink rate is the measure which shows the strongest results in the current study, which makes sense, echoing its use in many of the previous studies described above.

In conclusion, we showed that group size affected the blink rate in red and fallow deer groups in opposition to the group vigilance hypothesis, suggesting that social vigilance might be driving the blinking behaviour in these herds. Furthermore, we evaluated and discussed the validity and appropriateness of different measurement proxies for vigilance, and conclude that blink rate may provide a meaningful measure. While group vigilance is a well-studied area of animal behaviour, this technique provides the chance to re-examine past studies using a proxy that is less ambiguous in comparison to previous methods of used.

## ACKNOWLEDGMENTS

We thank three reviewers for their comments on a previous version of this study.

## DATA AVAILABILITY

The dataset and *R* code used to analyse it are freely available on *figshare* (Howard and Rands, 2025) at http://dx.doi.org/10.6084/m9.figshare.24931302

## REFERENCES

Allan, A.T.L., Hill, R.A., 2018. What have we been looking at? A call for consistency in studies of primate vigilance. Am J Phys Anthropol 165, 4–22. 10.1002/ajpa.23381

Beauchamp, G., 2019. On how risk and group size interact to influence vigilance. Biol Rev 94, 1918– 1934. 10.1111/brv.12540

Beauchamp, Guy, 2017. What can vigilance tell us about fear? Anim Sentience 2, 15.

Beauchamp, G., 2017. Half-blind to the risk of predation. Front Ecol Evol 5, 131. 10.3389/fevo.2017.00131

Bednekoff, P.A., Lima, S.L., 1998. Re–examining safety in numbers: interactions between risk dilution and collective detection depend upon predator targeting behaviour. Proc R Soc B 265, 2021– 2026. 10.1098/rspb.1998.0535

Blanchard, P., Pays, O., Fritz, H., 2017. Ticks or lions: trading between allogrooming and vigilance in maternal care. Anim Behav 129, 269–279. 10.1016/j.anbehav.2017.05.005

Cherry, R.L., Adair, H.S., Chen, T., Hendrix, D.V.H., Ward, D.A., 2020. Effect of attentional focus levels on spontaneous eyeblink rate in horses. Vet Opthalmol 23, 690–695. 10.1111/vop.12778

Childress, M.J., Lung, M.A., 2003. Predation risk, gender and the group size effect: does elk vigilance depend upon the behaviour of conspecifics? Anim Behav 66, 389–398. 10.1006/anbe.2003.2217

Creel, S., Winnie, J.A., Christianson, D., Liley, S., 2008. Time and space in general models of antipredator response: tests with wolves and elk. Anim Behav 76, 1139–1146. 10.1016/j.anbehav.2008.07.006

Cross, D.J., Marzluff, J.M., Palmquist, I., Minoshima, S., Shimizu, T., Miyaoka, R., 2013. Distinct neural circuits underlie assessment of a diversity of natural dangers by American crows. Proc R Soc B 280, 20131046. 10.1098/rspb.2013.1046

Crump, A., Arnott, G., Bethell, E.J., 2018. Affect-driven attention biases as animal welfare indicators: review and methods. Animals 8, 136. 10.3390/ani8080136

de la Peña, E., Pérez-González, J., Martín, J., Vedel, G., Carranza, J., 2021. The dark-ventral-patch of male red deer, a sexual signal that conveys the degree of involvement in rutting behavior. BMC Zool 6, 18. 10.1186/s40850-021-00083-9

Eccard, J.A., Meißner, J.K., Heurich, M., 2017. European roe deer increase vigilance when faced with immediate predation risk by Eurasian lynx. Ethology 123, 30–40. 10.1111/eth.12420

Favreau, F.-R., Goldizen, A.W., Pays, O., 2010. Interactions among social monitoring, anti-predator vigilance and group size in eastern grey kangaroos. Proc R Soc B 277, 2089–2095. 10.1098/rspb.2009.2337

Fernández-Juricic, E., Beauchamp, G., 2008. An experimental analysis of spatial position effects on foraging and vigilance in brown-headed cowbird flocks. Ethology 114, 105–114. 10.1111/j.1439-0310.2007.01433.x

Fernández-Juricic, E., Smith, R., Kacelnik, A., 2005. Increasing the costs of conspecific scanning in socially foraging starlings affects vigilance and foraging behaviour. Anim Behav 69, 73–81. 10.1016/j.anbehav.2004.01.019

Flamand, A., Robinet, L., Raskin, A., Braconnier, M., Bouhamidi, A., Derolez, G., Lochin, C., Helleu, C., Petit, O., 2025. The social dimension of equine welfare: social contact positively affects the emotional state of stalled horses. Anim Behav 221, 123055. 10.1016/j.anbehav.2024.123055

Fortin, D., Boyce, M.S., Merrill, E.H., Fryxell, J.M., 2004. Foraging costs of vigilance in large mammalian herbivores. Oikos 107, 172–180.

Friard, O., Gamba, M., 2016. BORIS: a free, versatile open-source event-logging software for video/audio coding and live observations. Methods Ecol Evol 7, 1325–1330. 10.1111/2041-210X.12584

Ginkel, H.A.L. van, Smit, C., Kuijper, D.P.J., 2019. Behavioral response of naïve and non-naïve deer to wolf urine. PLoS One 14, e0223248. 10.1371/journal.pone.0223248

Hoppe, D., Helfmann, S., Rothkopf, C.A., 2018. Humans quickly learn to blink strategically in response to environmental task demands. Proc Natl Acad Sci USA 115, 2246–2251. 10.1073/pnas.1714220115

Howard, T.R., Rands, S.A., 2025. Supplementary Material for “Video-recorded measures of blinking differ with group size in red deer Cervus elaphus and fallow deer Dama dama.” Figshare 10.6084/m9.figshare.24931302

Hoyle, Z.E., Miller, R.A., Rands, S.A., 2021. Behavioural synchrony between fallow deer Dama dama is related to spatial proximity. BMC Ecol Evo 21, 79. 10.1186/s12862-021-01814-9

Hurlbert, S.H., 1984. Pseudoreplication and the design of ecological field experiments. Ecol Monogr 54, 187–211.

Jayakody, S., Sibbald, A.M., Gordon, I.J., Lambin, X., 2008. Red deer Cervus elephus vigilance behaviour differs with habitat and type of human disturbance. Wildl Biol 14, 81–91. 10.2981/0909-6396(2008)14[81:RDCEVB]2.0.CO;2

Kjær, L.J., Schauber, E.M., Nielsen, C.K., 2008. Spatial and temporal analysis of contact rates in female white-tailed deer. J Wildl Manage 72, 1819–1825. 10.2193/2007-489

Lehtonen, J., Jaatinen, K., 2016. Safety in numbers: the dilution effect and other drivers of group life in the face of danger. Behav Ecol Sociobiol 70, 449–458. 10.1007/s00265-016-2075-5

Lelláková, M., Pavlak, Alexander, Lešková, Lenka, Florián, Martin, Skurková, Lenka, Mesarčová, Lýdia, Kottferová, Lucia, Takáčová, Daniela, and Kottferová, J., 2023. Monitoring blinks and eyelid twitches in horses to assess stress during the samples collection process. J Appl Anim Welfare Sci 26, 530–539. 10.1080/10888705.2021.2008249

Li, Z., Jiang, Z., 2008. Group size effect on vigilance: evidence from Tibetan gazelle in Upper Buha River, Qinghai-Tibet Plateau. Behav Process 78, 25–28. 10.1016/j.beproc.2007.11.011

Li, Z., Jiang, Z., Beauchamp, G., 2009. Vigilance in Przewalski’s gazelle: effects of sex, predation risk and group size. J Zool 277, 302–308. 10.1111/j.1469-7998.2008.00541.x

Matsumoto-Oda, A., Okamoto, K., Takahashi, K., Ohira, H., 2018. Group size effects on inter-blink interval as an indicator of antipredator vigilance in wild baboons. Sci Rep 8, 10062. 10.1038/s41598-018-28174-7

McSweeney, F.K., Swindell, S., 2002. Common processes may contribute to extinction and habituation. J Gen Psychol 129, 364–400. 10.1080/00221300209602103

Merkies, K., Ready, C., Farkas, L., Hodder, A., 2019. Eye blink rates and eyelid twitches as a non-invasive measure of stress in the domestic horse. Animals 9, 562. 10.3390/ani9080562

Pecorella, I., Fattorini, N., Macchi, E., Ferretti, F., 2019. Sex/age differences in foraging, vigilance and alertness in a social herbivore. Acta Ethol 22, 1–8. 10.1007/s10211-018-0300-0

Pecorella, I., Ferretti, F., Sforzi, A., Macchi, E., Pecorella, I., Ferretti, F., Sforzi, A., Macchi, E., 2016. Effects of culling on vigilance behaviour and endogenous stress response of female fallow deer. Wildl Res 43, 189–196. 10.1071/WR15118

R Development Core Team, 2023. R: a language and environment for statistical computing. R Foundation for Statistical Computing, Vienna.

Rands, S.A., 2021. Phylogenetically-controlled correlates of primate blinking behaviour. PeerJ 9, e10950. 10.7717/peerj.10950

Rands, S.A., 2015. Nearest-neighbour clusters as a novel technique for assessing group associations. R Soc Open Sci 2, 140232. 10.1098/rsos.140232

Rands, S.A., Muir, H., Terry, N.L., 2014. Red deer synchronise their activity with close neighbours. PeerJ 2, e344. 10.7717/peerj.344

Ranti, C., Jones, W., Klin, A., Shultz, S., 2020. Blink rate patterns provide a reliable measure of individual engagement with scene content. Sci Rep 10, 8267. 10.1038/s41598-020-64999-x

Rowe, Z.W., Robins, J.H., Rands, S.A., 2023. Red deer Cervus elaphus blink more in larger groups. Ecol Evol 13, e9908. 10.1002/ece3.9908

Sansom, A., Cresswell, W., Minderman, J., Lind, J., 2008. Vigilance benefits and competition costs in groups: do individual redshanks gain an overall foraging benefit? Anim Behav 75, 1869–1875. 10.1016/j.anbehav.2007.11.005

Sibbald, A.M., Hooper, R.J., McLeod, J.E., Gordon, I.J., 2011. Responses of red deer (Cervus elaphus) to regular disturbance by hill walkers. Eur J Wildl Res 57, 817–825. 10.1007/s10344-011-0493-2

Tada, H., Omori, Y., Hirokawa, K., Ohira, H., Tomonaga, M., 2013. Eye-blink behaviors in 71 species of primates. PLoS One 8, e66018. 10.1371/journal.pone.0066018

Tomberg, C., Petagna, M., de Selliers de Moranville, L.-A., 2024. Spontaneous eye blinks in horses (Equus caballus) are modulated by attention. Sci Rep 14, 19336. 10.1038/s41598-024-70141-y

Welp, T., Rushen, J., Kramer, D.L., Festa-Bianchet, M., de Passillé, A.M.B., 2004. Vigilance as a measure of fear in dairy cattle. Appl Anim Behav Sci 87, 1–13. 10.1016/j.applanim.2003.12.013

Wickham, H., 2009. ggplot2: elegant graphics for data analysis. Springer-Verlag, New York.

Yorzinski, J.L., 2020. Blinking behavior in great-tailed grackles (Quiscalus mexicanus) increases during simulated rainfall. Ethology 126, 519–527. 10.1111/eth.13003

Yorzinski, J.L., 2016. Eye blinking in an avian species is associated with gaze shifts. Sci Rep 6, 32471. 10.1038/srep32471

Yorzinski, J.L., Argubright, S., 2019. Wind increases blinking behavior in great-tailed grackles (Quiscalus mexicanus). Front Ecol Evol 7, 330. 10.3389/fevo.2019.00330

Yorzinski, J.L., Walker, M.K., Cavalier, R., 2021. A songbird strategically modifies its blinking behavior when viewing human faces. Anim Cogn 24, 787–801. 10.1007/s10071-021-01476-6

Zheng, W., Beauchamp, G., Jiang, X., Li, Z., Yang, Q., 2013. Determinants of vigilance in a reintroduced population of Père David’s deer. Curr Zool 59, 265–270. 10.1093/czoolo/59.2.265

